# MOSQUITO MICROBIOMES OF RWANDA: CHARACTERIZING MOSQUITO HOST AND MICROBIAL COMMUNITIES IN THE LAND OF A THOUSAND HILLS

**DOI:** 10.1101/2022.08.03.502589

**Authors:** Amanda G. Tokash-Peters, Jean Damascene Niyonzima, Mirielle Kayirangwa, Simon Muhayimana, Ivan W. Tokash, Jaimy D. Jabon, Sergio G. Lopez, Douglas C. Woodhams

**Affiliations:** College of Science and Mathematics, University of Massachusetts Boston, Boston, MA; Center of Excellence in Biodiversity, University of Rwanda, Huye, Rwanda

**Keywords:** Rwanda, mosquito microbiome, vector-borne disease, disease ecology

## Abstract

Mosquitoes are a complex nuisance around the world, and tropical countries bear the greatest brunt of the burden of mosquito-borne diseases. Rwanda has had success in reducing malaria and some arboviral diseases over the last few years, but still faces challenges to elimination. By building our understanding of *in situ* mosquito communities in Rwanda at a disturbed, human-occupied site and at a natural, preserved site, we can build our understanding of natural mosquito microbiomes toward the goal of implementing novel microbial control methods. Here, we examined the composition of collected mosquitoes and their microbiomes at two diverse sites using Cytochrome c Oxidase I sequencing and 16S V4 barcode sequencing. The majority of mosquitoes captured and characterized in this study are the first-known record of their species for Rwanda but have been characterized in other nations in East Africa. Beta diversity metrics were significantly different between sampling sites, mosquito genera, and mosquito species, but not between mosquito sexes, catch method, or presumed bloodfed status. Bacteria of interest for arbovirus control, *Asaia, Serratia*, and *Wolbachia*, were found in abundance at both sites, but were more prevalent at the disturbed site and varied greatly by species. Additional studies to build our understanding of naturally-formed microbial communities are essential to safely employing microbial control methods and further reducing the burden of mosquito-borne diseases.

## Introduction

Mosquito-borne diseases disproportionately impact nations in the tropics, with Sub-Saharan Africa bearing a sizeable burden (World Health Organization, 2017; World Health Organization Africa Region, 2019). Annually around the world, there are between 100-400 million cases of Dengue fever, 200,000 cases of yellow fever, 228 million cases of malaria, and several hundred periodic cases of Rift Valley fever (Bhatt et al., 2013; Brady et al., 2015; CDC, 2018; Garske et al., 2014; Nanyingi et al., 2015; Shearer et al., 2018; World Health Organization, 2017, 2018a, 2018b). While some control efforts have stagnated over time, Rwanda has effectively reduced the number of malaria cases by 430,000 from 2016 to 2017 (Ingabire et al., 2017; World Health Organization, 2018b).

Despite this remarkable reduction, challenges still exist in further reducing arbovirus and parasite transmission. One mode of control that shows potential in pushing toward reduction goals is microbial manipulation of mosquito hosts (Bian et al., 2013; Cirimotich et al., 2011; Dodson et al., 2017; Hoffmann et al., 2011; McMeniman et al., 2009; Moreira et al., 2009; Walker et al., 2011; S. Wang et al., 2017; Weiss & Aksoy, 2011). Introduction of bacteria of the genera *Asaia, Serratia*, and *Wolbachia* into the mosquito microbiome have all been implicated as potential methods for pathogen and parasite control. These bacteria can either trigger an immune response in the mosquito that can help reduce prevalence of pathogens and parasites or can directly compete with invading microbes (Hegde et al., 2018; Jupatanakul et al., 2014; Rancès et al., 2012; Sigle et al., 2016; Y.-H. Wang et al., 2017). Though the World Mosquito Program has begun implementing the release of *Wolbachia*-infected *Aedes* mosquitoes with success, there are still numerous uncertainties regarding *in situ* release and efficacy of these methods (Hughes et al., 2012; Murray et al., 2016; O’Neill, 2018; Zélé et al., 2014).

In order to better understand the probability of success for microbial methods of control in East Africa, background understanding of naturally-formed mosquito host and microbiome communities must first be formed. While we are still building our understanding of factors that influence the formation of mosquito microbiomes, we know that the environment plays an important role (Alto et al., 2008; Caporaso et al., 2011; Dickson et al., 2017; Y. Wang et al., 2011a).

In this study, we aimed to characterize the mosquito and microbial communities of two diverse sites in Rwanda to help elucidate factors influencing microbiome assemblage. We hypothesized that there would be differences in microbial composition across the sampling locations, mosquito taxa, mosquito sexes, and whether mosquito females were visibly bloodfed or not. This study also aimed to describe mosquito communities at both sampling locations and help to establish potentially novel records for mosquitoes in Rwanda.

## Methods

### Site Selection and Research Permit

Two sampling sites in Rwanda were selected for comparison in the characterization of mosquito microbiomes. Samples were collected in February 2019 between the peak dry and wet seasons. Sites were selected that had nearby (<500m) frequently traversed walking paths. Sites were also secondary growth forests with similar canopy and ground cover. The first selected was near Kigali, upland of Nyabugogo marsh, and in a grove of trees in a highly disturbed area with densely packed temporary housing with poor sanitation conditions. The second site selected was the arboretum on the Huye campus of the University of Rwanda which contains a footpath frequently traversed by members of the community but no settlements in the immediate vicinity of the site.

### Ethics Statement

Field sampling was performed under the authority of the Rwandan National Council for Science and Technology (NCST) under Permit # NCST/482/62/2018.

### Specimen Collection, Storage, and Extraction

Mosquito samples were collected using human landing catch (HLC) prior to, during, and for three hours following sunrise using either aspirator or non-destructive hand-grab collection. For specimens caught with hands, 70% ethanol sanitizer was used in between grabs. Additional samples at each site were collected while resting on surrounding foliage using either aspirator or hand catch. Samples were frozen on collection at −20°C until they could be processed, and then had the head and legs removed before being homogenized. Samples were individually stored in 1X DNA RNA Shield (Zymo Research) and were extracted using Quick DNA/RNA MagBead kits (Zymo Research) and stored at −80°C.

### Cytochrome C Oxidase Subunit I Barcoding

Mosquito species identification was performed using cytochrome c oxidase subunit I conserved gene barcoding. PCR was performed using the standard LCO1490 and HCO2198 primers to target the 710-bp fragment (Folmer et al., 1994). Polymerase chain reaction (PCR) was executed in duplicate using 5Prime HotMasterMix (Quantabio) (10ul), Milli-Q Ultrapure Water (11ul), 10uM LCO1490 (1uL), 10uM HCO2198 (1uL), and template DNA (2uL) (Folmer et al., 1994). The PCR cycling conditions were 94°C for 3 minutes, 35 cycles of 94°C for 45 seconds, 50°C for 60 seconds, and 72°C for 90 seconds, followed by 72°C for 10 minutes with a holding temperature of 4°C. Samples were then pooled and sent to MC Lab (South San Francisco, CA) for PCR clean-up and sequencing using an ABI 3730XL sequencer. Upon retrieval of sequences from MC Lab, FASTA sequences were matched to mosquito species-level identifications using the BOLD Systems Identification Engine for all records on BOLD as of March 2020 (Ratnasingham & Hebert, 2007). The highest percentage match at the species level of identification was used as the positive identification of the sample. Samples with exceptionally poor species level matches (<85%) were not included in further analyses utilizing the taxonomic identification. To ensure that identification of each species would be highly likely in Rwanda, species were all cross-referenced in BOLD, the Walter Reed Biosystematics Unit Systematic Catalogue of Culicidae, and the broader literature for known records of presence in East Africa. Identified species of mosquitoes were broadly classified into generalized habitat use groups (Domestic, Sylvatic, Ubiquitous, Unknown, or Wetland) based on existing literature of habitat use, particularly in East Africa whenever possible.

### Microbiome Sequencing and Preparation

The 16S rRNA V4 gene region was amplified by PCR using the standard 515F and 806R primers with barcodes on the forward primer consistent with the Earth Microbiome Project (EMP) protocols (Apprill et al., 2015; Caporaso et al., 2011, 2012; Parada et al., 2016; Walters et al., 2016). PCR was performed with minor modifications to the EMP protocols, amplifying samples and controls in duplicate using 5Prime HotMasterMix (Quantabio) (10ul), Milli-Q Ultrapure Water (8.5ul), 10uM 806R (3.5uL), 10uM barcoded 515F (1uL), and template DNA (2uL) for a total of 25ul per well. Cycling conditions followed the standard EMP protocol and can also be found listed above in the methods section on COI barcoding. PCR-amplified samples were then normalized using the Mag-Bind Pure Library Normalization Kit (Omega Bio-Tek) and pooled for library quantification and subsequent dilution using a Qubit 2 Fluorometer (Invitrogen). PhiX was added according to recommendations from the Earth Microbiome Project and the library was sequenced using an Illumina MiSeq v2 300-cycle kit (*The Earth Microbiome Project-Protocols and Standards*, 2020; Thompson et al., 2017).

### Bioinformatic and Statistical Analyses

Analysis of mosquito microbiomes was undertaken using QIIME2 and the vegan, MicrobeR, and qiime2r packages in R. All related code files, the mapping file, and the demultiplexed sequences can found in Appendices I and II. Microbiome data were demultiplexed, deblurred, quality filtered, and rarefied to 2000 sequences per sample in QIIME2. Kruskal-Wallis pairwise tests were performed to analyze differences in sOTU richness between experimental groups. PERMANOVAs were performed to assess statistical differences in the beta diversity of microbial communities between experimental groups using the permanova function in QIIME2. All figures were generated in R or Excel.

## Results

### Mosquito Community Results

Between both the Nyabugogo and Huye Campus sites, 447 individuals belonging to forty species of mosquitoes were successfully identified across six genera. Female captures greatly outnumbered male captures, at approximately double the rate across both sampling sites. Based on BOLD and the Walter Reed Biosystematics Unit Systematic Catalogue of Culicidae, 36 of the 40 species collected were potentially the first record of these mosquitoes in Rwanda, though nearly all of these species were found in other east African nations. Culex were the most prevalent genus of mosquitoes at both sites. Overall, *Lutzia tigripes, Culex univitattus,* and *Culex decens* were the most frequently captured individuals. At the Huye Campus site, *Culex univitattus, Culex decens,* and *Aedes mcintoshi* were the most frequently encountered species, while *Lutzia tigripes, Culex univitattus,* and *Culex striatipes* were the most encountered at Nyabugogo.

The community composition between the sites differed substantially (*Fig. 1*). The Huye Campus site had greater numbers of Aedes, Coquilletidia, and Eretmapodites, whereas Nyabugogo had more Lutzia and accounted for the single Anopheles mosquito collected. Culex mosquitoes were the majority of mosquitoes at both sites, though the species composition of Culex differed.

**Figure 1.**
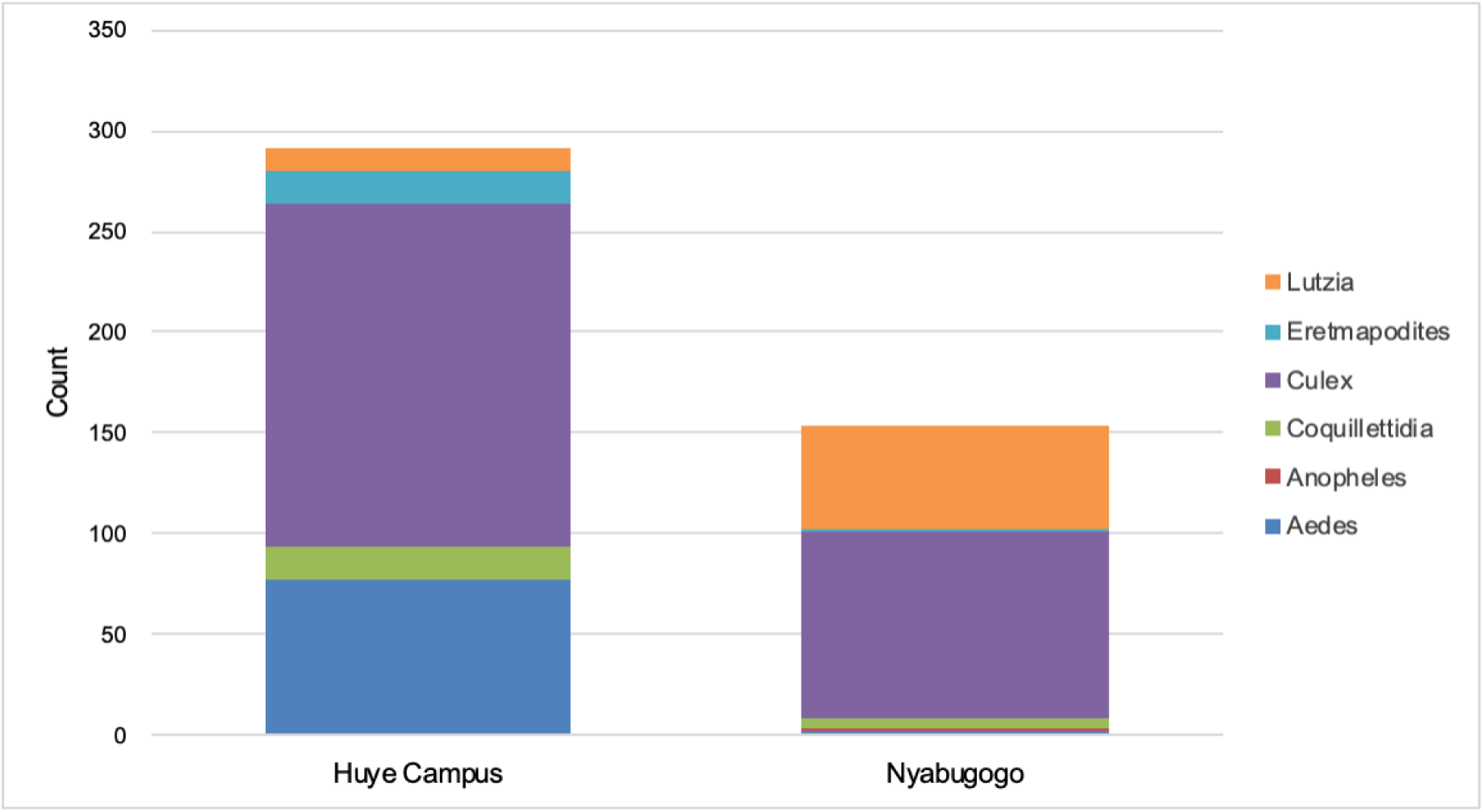
Stacked barchart comparing the relative count and assemblage of mosquitoes captured at the Huye Campus (arboretum) and Nyabugogo (highly disturbed) sites. *Culex* mosquitoes were the majority sampled at both sites. *Aedes* mosquitoes were highly prevalent at the Huye Campus site and *Lutzia* mosquitoes were common at Nyabugogo.

In terms of the generalized habitat use assignments, mosquitoes classified as ubiquitous were the most abundant overall, as well as at the Huye Campus site (*Fig. 2*). At the Nyabugogo site, domestic mosquitoes were most common and nearly double the number of ubiquitous mosquitoes captured. Sylvatic mosquitoes were highly abundant at the Huye Campus site, while they were far less prevalent at the Nyabugogo site. Specifically in terms of relative abundance, there were more wetland mosquitoes at the Huye Campus site than Nyabugogo.

**Figure 2.**
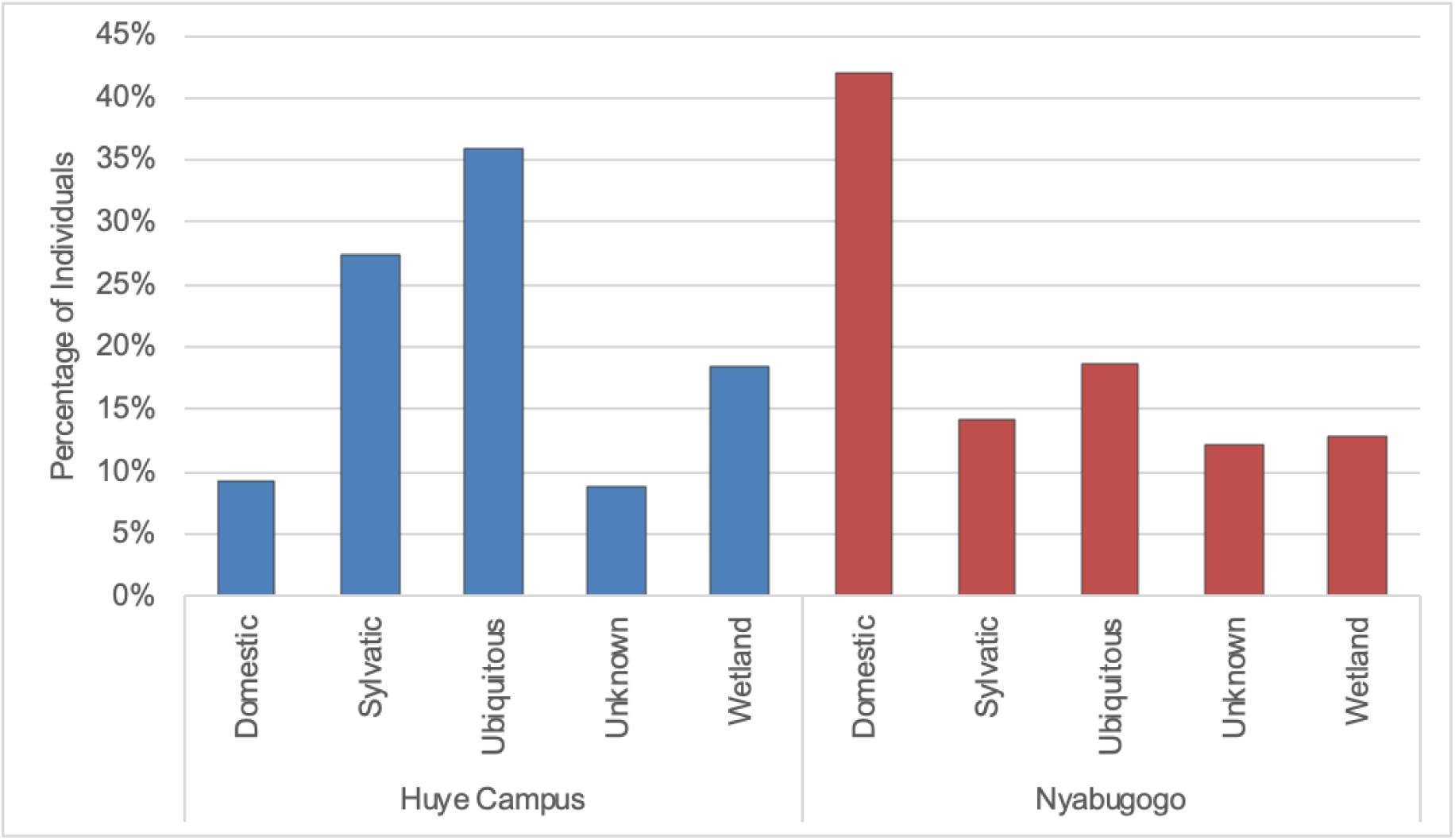
Relative abundance in percentage of all mosquitoes by sampling site for each generalized habitat use categorization. Sylvatic and wetland mosquitoes were more common at the Huye Campus site, while domestic mosquitoes were more abundant at the Nyabugogo site (Burke et al., 2010; Chabaud & Ovazza, 1958; Diallo et al., 2012; Gibbins, 1942; A. J. Haddow, 1956; Junglen et al., 2009; Lu et al., 2013; Lutomiah et al., 2013; Miller et al., 2000; Mutebi et al., 2013a; Muturi et al., 2006, 2007; Ndiaye et al., 2016a; Obame-Nkoghe et al., 2017; Robert et al., 1998; Roiz et al., 2015; Sallam et al., 2013; Self et al., 1973; SNOW, 1987; Soti et al., 2012; Zahouli et al., 2017).

### Microbiome Diversity

#### Alpha Diversity of the Microbiome: sOTU Richness

Visibly blood fed and non-blood fed mosquitoes were marginally different in terms of sOTU richness of the microbiome in a pairwise Kruskal-Wallis test using observed OTUs (p= 0.06). There was no other significant difference between any groups in terms of microbiome sOTU richness, including sex, location, catch method, collection date, or processing date.

#### Alpha Diversity of the Microbiome: Shannon Diversity Index

When using the Shannon Diversity Index of the microbiome to run Kruskal-Wallis grouped and pairwise tests, there was no significant difference between locations, sexes, genera, species, catch methods, or visible bloodfeeding status.

#### Beta Diversity of the Microbiome: Jaccard Distances

PERMANOVAs were run to analyze differences in microbiome beta diversity between key groups. Each PERMANOVA was run with 999 permutations. There were significant differences when analyzing groups using Jaccard distances between genera (n=392, pseudo-F statistic= 1.44492, p= 0.001), species (n=393, pseudo-F statistic = 1.15976, p= 0.001), and location (n= 466, pseudo-F statistic = 2.29368, p= 0.001), but not whether the mosquito was visibly blood-fed (n= 464, pseudo-F statistic = 1.04428, p= 0.258).

#### Beta Diversity of the Microbiome: Weighted Unifrac Distances

There were significant differences when tested using PERMANOVAs between weighted Unifrac values for sampling locations (n=442, pseudo-F statistic = 9.02676, p=0.001), genera (n=466, pseudo-F statistic =2.48844, p=0.002), species (n=466, pseudo-F statistic = 2.01142, p=0.001), catch dates (n= 466, pseudo-F statistic = 5.16716, p=0.001), and processing dates (n=466, pseudo-F statistic = 2.97262, p=0.001). As only one site was sampled per day and which groups of mosquitoes were processed on which date was non-random, these were considered to be an artifact of these variables strongly co-varying with location. Mosquito sexes (n=454, pseudo-F statistic = 2.02918, p=0.068), visibly bloodfed status (n= 466, pseudo-F statistic = 0.964349, p=0.457), and catch methods (n= 466, pseudo-F statistic = 1.53627, p=0.115) were all not significantly different. The principal component analysis in Figure 4 demonstrates the differences in community composition of the mosquito genera at both sampling locations. Aedes and Coquillettidia from the Huye Campus site have high dispersion in their microbial community compositions compared to those genera at Nyabugogo. Lutzia has higher dispersion at Nyabugogo, while Culex is fairly disperse at both sites. A principal coordinate analysis of species level variation in microbiome communities can be found in the supplement (*Fig. S1*).

**Figure 3.**
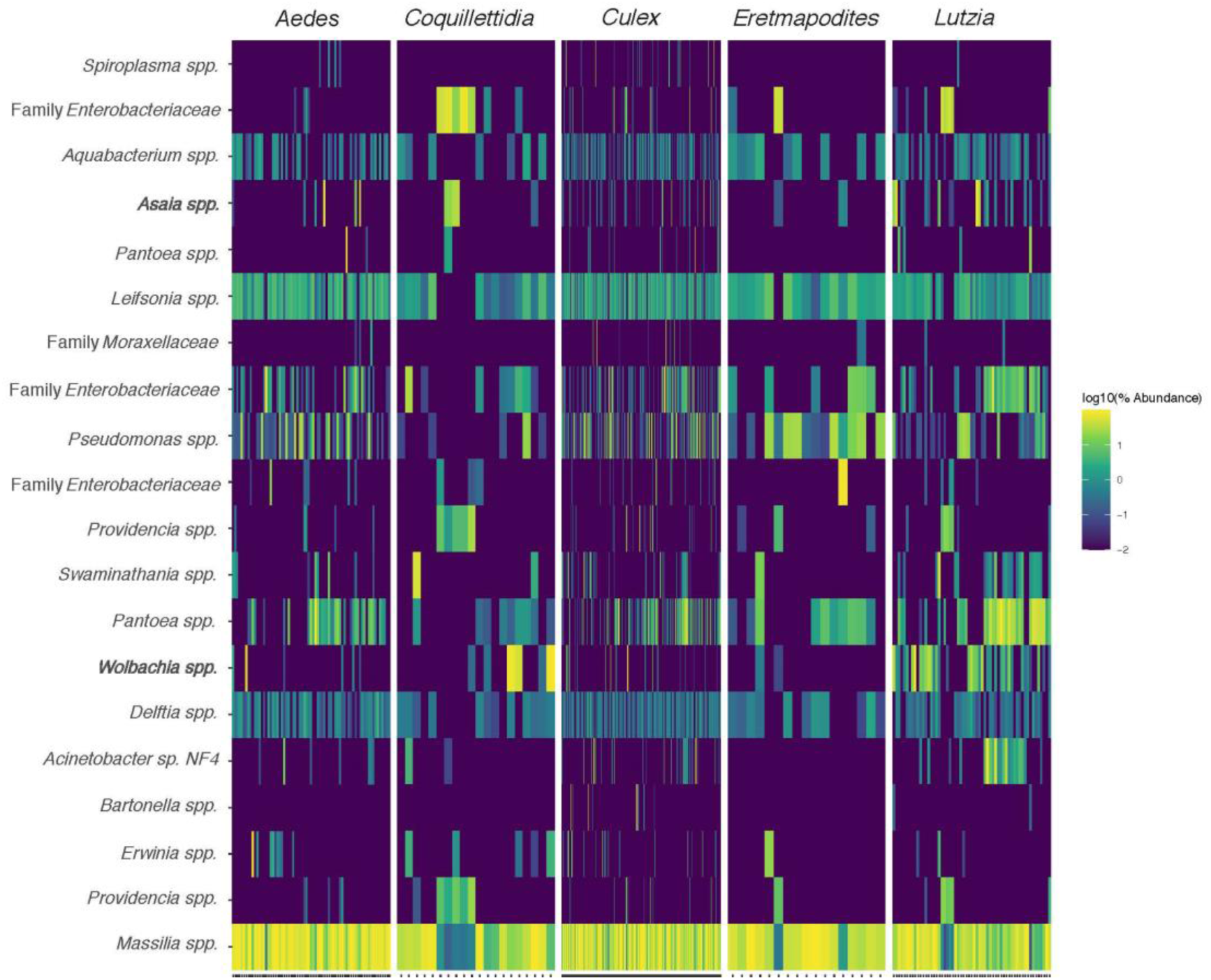
Heatmap of the 20 most abundant sOTUs across all mosquito samples, grouped by genus. *Massilia spp.* was the most overall abundant sOTU, with high abundances of *Leifsonia spp*., *Delftia spp., Pantoea spp*., and *Pseudomonas spp*. Other notable members of the core microbiome include *Wolbachia spp*. and *Asaia spp*. that have been noted for their role in the control of arboviral transmission.

**Figure 4.**
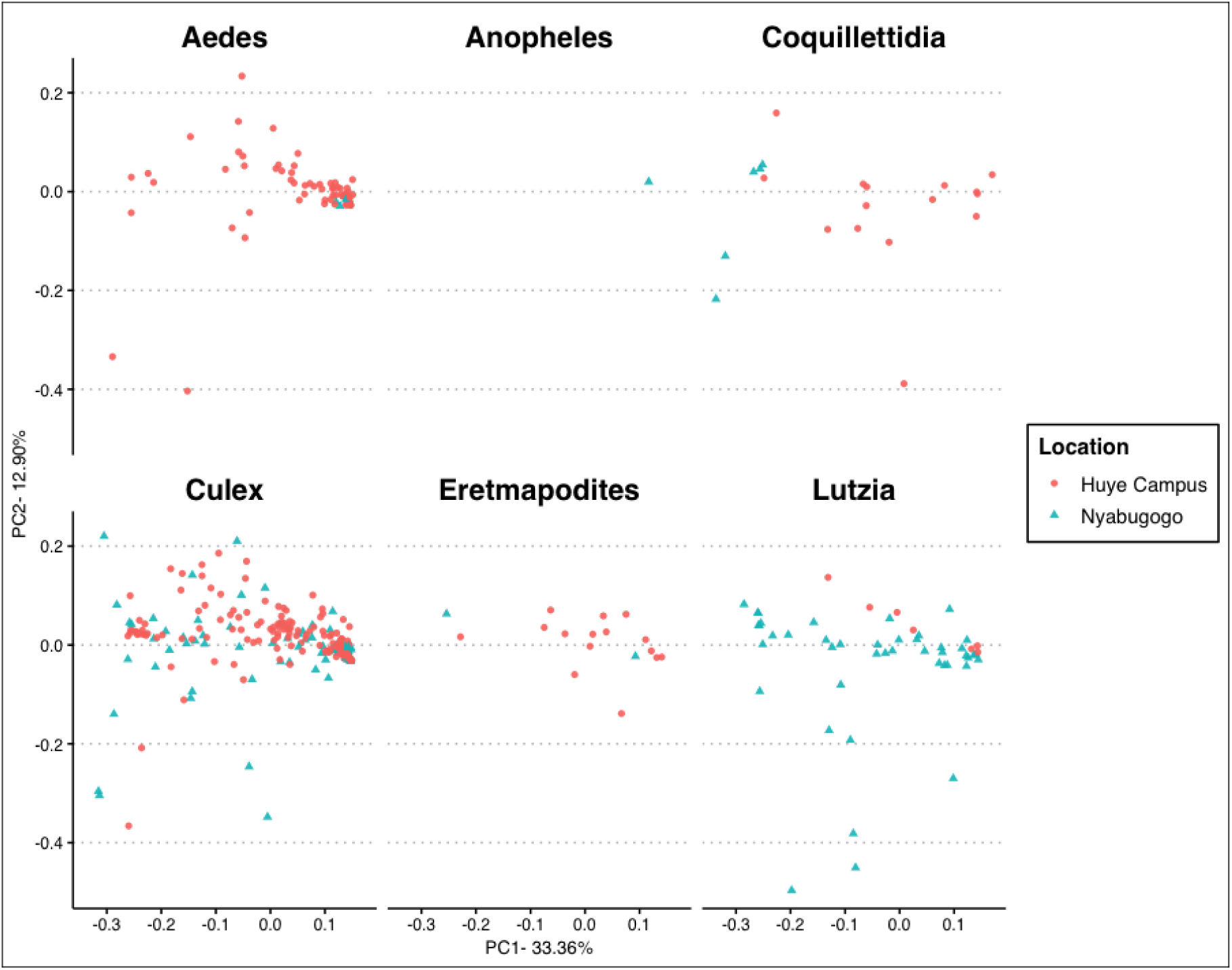
Principle coordinate analysis made with weighted Unifrac distances to demonstrate the relative community composition of the microbiome for each genus by sampling location. *Aedes* and *Coquillettidia* mosquitoes are disperse at the Huye Campus site, while *Lutzia* is most disperse at the Nyabugogo site. *Culex* assemblages are disperse at both sites.

### The Core Microbiome

Core members of the mosquito microbiome were considered to be the top 20 most abundant sOTUs across all mosquito samples (*Fig.3*). The most abundant sOTUs were *Massillia spp., Leifsonia spp.,* and *Delftia spp.* Other noteable members of the core microbiome include *Wolbachia spp.* and *Asaia spp.,* as these microbes have been implicated as having a role in the transmission or control of arboviruses.

### Wolbachia and Bacteria of Anti-arboviral Interest

*Wolbachia, Asaia*, and *Serratia* are genera of bacteria that have been implicated in controlling transmission of some arboviruses and parasites (Alfano et al., 2019; Cirimotich et al., 2011; Moreira et al., 2009; Novakova et al., 2017; S. Wang et al., 2017). All *Coquillettidia* species caught in Rwanda were strong carriers of *Wolbachia* (*Fig. 5). Aedes, Culex, Eretmapodites*, and *Lutzia* all had some species that carried *Wolbachia*.

**Figure 5.**
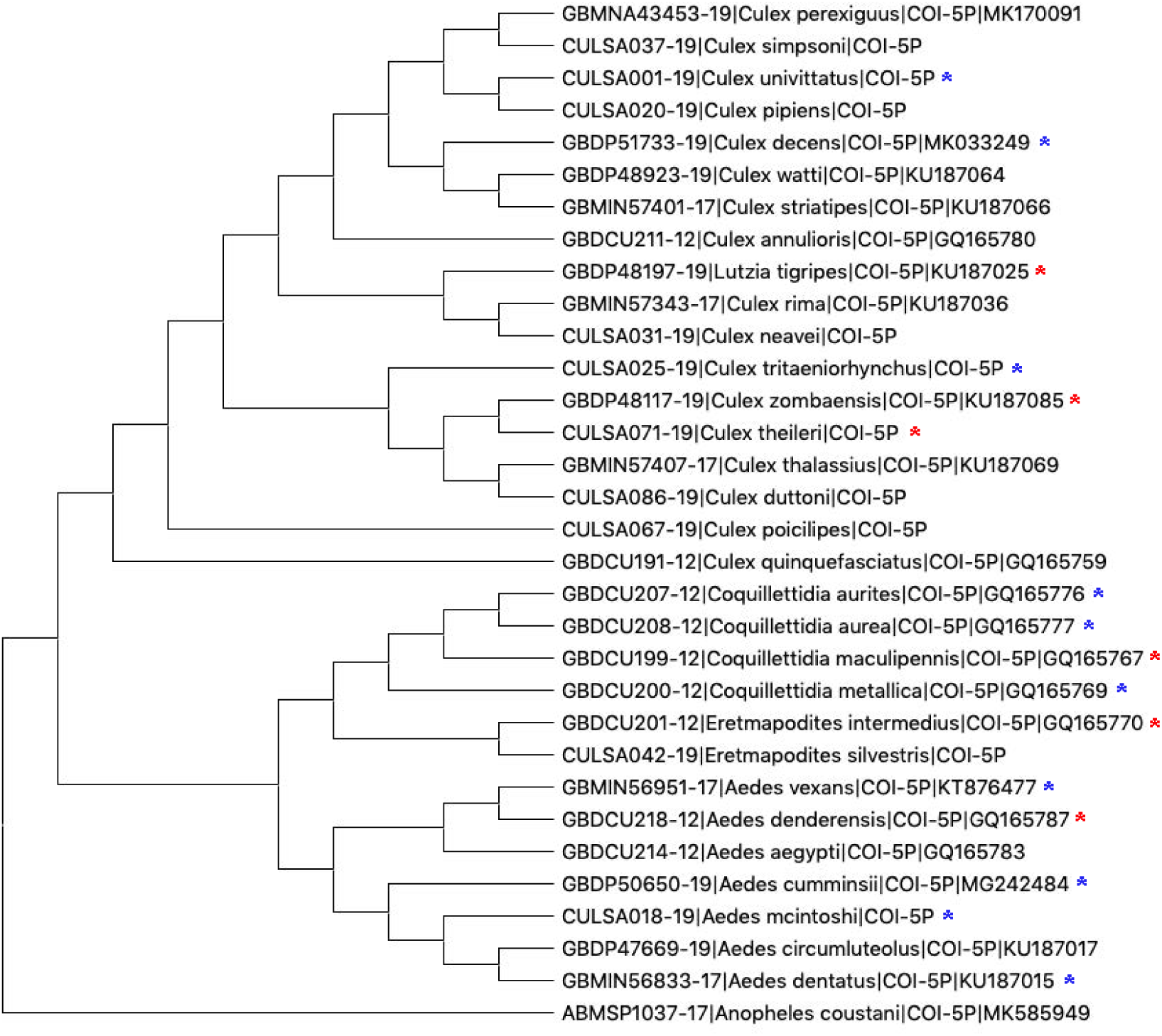
Phylogenetic tree of mosquito species found at Rwandan sampling sites created with publicly available high-quality and long-length cytochrome c oxidase II (COI) sequences from BOLD derived from primarily Ugandan, Kenyan, and South African mosquitoes. Sequences were aligned and the tree was created using the neighbor-joining method with 100 Maximum Likelihood bootstrap replicates in MEGA X (Kumar et al., 2018). Asterisks denote Rwandan species captured in this project that carried *Wolbachia spp.*, with the blue asterisks indicating between 0.5-100 average sequences and red asterisks indicating >100 average sequences.

The average number of sequences for each of the bacterial genera of interest varied widely by mosquito host species. Overall, the Nyabugogo site, which was highly disturbed and close to human habitation, had higher numbers of average sequences of *Asaia, Serratia*, and *Wolbachia* than the preserved and heavily-forested Huye Campus site (Table 1). Interestingly, some mosquito host species that were found in abundance at both sampling locations had considerably different average numbers of sequences. For example, *Lutzia tigripes* was more relatively abundant at Nyabugogo and also had substantially higher average sequences of *Asaia* and *Wolbachia* at that same site. *Culex striatipes* (domestic) and *Culex rima* (sylvatic), which were also more abundant at Nyabugogo, had substantially higher average *Asaia* and *Serratia* sequences at the Nyabugogo site compared to the Huye Campus as well. *Culex theileri,* a sylvatic mosquito, also had higher average sequences of *Serratia* and *Wolbachia* at Nyabugogo. In contrast, *Aedes mcintoshi* (ubiquitous) was found at both sites and was much more abundant at the Huye Campus site, and had higher average sequences of *Asaia* and *Serratia* (but not *Wolbachia*) at the Huye Campus site.

**Table 1.** Average number of sequences for mosquito microbiome members of arboviral control interest (Asaia, Serratia, and Wolbachia). Mosquito species are separated by sampling location and are designated with their general habitat use classification. These average sequence values are based on samples rarefied to 2000 sequences. Nyabugogo overall had higher average numbers of Asaia, Serratia, and Wolbachia than the Huye Campus site.

| Mosquitoes by Location | Generalized Habitat Use | Average Asaia Seqs/Sample | Average Serratia Seqs/Sample | Average Wolbachia Seqs/Sample |
| --- | --- | --- | --- | --- |
| Huye Campus |  | 17.1 | 4.2 | 49.9 |
| Culex duttoni | Domestic | 0.0 | 4.5 | 0.0 |
| Culex striatipes | Domestic | 0.0 | 0.5 | 0.0 |
| Culex thalassius | Domestic | 0.0 | 3.5 | 0.0 |
| Lutzia tigripes | Domestic | 3.5 | 2.9 | 2.0 |
| Aedes argenteopunctatus | Sylvatic | 0.0 | 0.0 | 0.0 |
| Aedes cumminsii | Sylvatic | 0.0 | 0.0 | 0.5 |
| Aedes denderensis | Sylvatic | 486.0 | 7.5 | 773.0 |
| Aedes dentatus | Sylvatic | 12.5 | 5.5 | 0.8 |
| Aedes ingrami | Sylvatic | 0.0 | 0.0 | 0.0 |
| Coquillettidia aurea | Sylvatic | 0.3 | 3.7 | 96.5 |
| Coquillettidia maculipennis | Sylvatic | 87.8 | 0.3 | 821.0 |
| Culex neavei | Sylvatic | 0.0 | 0.5 | 0.0 |
| Culex rima | Sylvatic | 0.0 | 3.0 | 0.0 |
| Culex simpsoni | Sylvatic | 9.3 | 1.5 | 0.0 |
| Culex theileri | Sylvatic | 0.0 | 2.4 | 201.7 |
| Culex watti | Sylvatic | 0.0 | 14.0 | 0.0 |
| Eretmapodites intermedius | Sylvatic | 0.5 | 3.5 | 163.8 |
| Eretmapodites silvestris | Sylvatic | 0.0 | 2.5 | 0.0 |
| Aedes circumluteolus | Ubiquitous | 0.0 | 0.0 | 0.0 |
| Aedes mcintoshi | Ubiquitous | 40.3 | 5.5 | 13.1 |
| Aedes vexans | Ubiquitous | 0.0 | 10.4 | 0.0 |
| Culex annulioris | Ubiquitous | 158.5 | 0.0 | 0.0 |
| Culex decens | Ubiquitous | 3.1 | 3.4 | 0.7 |
| Culex pipiens | Ubiquitous | 0.0 | 3.0 | 0.0 |
| Culex zombaensis | Ubiquitous | 0.0 | 1.0 | 730.5 |
| Culex mali sp. 4 | Unknown | 0.0 | 4.6 | 30.7 |
| Culex mali sp.1 | Unknown | 64.0 | 0.0 | 0.0 |
| Culex sp. GP-B | Unknown | 0.0 | 1.9 | 104.6 |
| Culex sp. M63YA | Unknown | 0.6 | 1.7 | 1.7 |
| Culex perexiguus | Wetland | 0.0 | 3.0 | 0.0 |
| Culex univittatus | Wetland | 4.1 | 6.8 | 0.5 |
| Nyabugogo |  | 58.1 | 13.5 | 74.9 |
| Culex striatipes | Domestic | 121.8 | 151.1 | 0.0 |
| Lutzia tigripes | Domestic | 97.0 | 3.2 | 172.2 |
| Coquillettidia aurites | Sylvatic | 2.0 | 0.0 | 1.0 |
| Coquillettidia metallica | Sylvatic | 262.3 | 0.0 | 1.5 |
| Culex neavei | Sylvatic | 1.1 | 8.0 | 0.0 |
| Culex rima | Sylvatic | 6.8 | 2.8 | 0.0 |
| Culex theileri | Sylvatic | 0.0 | 21.0 | 370.0 |
| Eretmapodites intermedius | Sylvatic | 0.5 | 0.5 | 247.5 |
| Aedes mcintoshi | Ubiquitous | 0.0 | 4.5 | 4.0 |
| Aedes vexans | Ubiquitous | 3.0 | 1.0 | 59.0 |
| Anopheles coustani | Ubiquitous | 0.0 | 2.0 | 0.0 |
| Culex decens | Ubiquitous | 0.1 | 6.3 | 7.7 |
| Culex pipiens | Ubiquitous | 27.0 | 2.0 | 0.0 |
| Culex poicillipes | Ubiquitous | 876.0 | 0.0 | 0.0 |
| Culex zombaensis | Ubiquitous | 0.0 | 15.3 | 228.0 |
| Culex mali sp. 4 | Unknown | 6.5 | 11.5 | 0.0 |
| Culex sp. GP-B | Unknown | 14.6 | 11.2 | 0.0 |
| Culex sp. M63YA | Unknown | 3.5 | 2.1 | 0.0 |
| Culex sp. T15 | Unknown | 0.0 | 0.0 | 0.0 |
| Culex tritaeniorhynchus | Wetland | 96.0 | 0.0 | 17.0 |
| Culex univittatus | Wetland | 3.9 | 3.0 | 0.9 |
| Average Sequences/2000 Across All Samples |  | 31.0 | 7.4 | 58.4 |

## Discussion

Rwandan mosquito microbiomes remain understudied and have the potential to provide imperative information in the regional fight against arboviral and *Plasmodium* infections. Here, we present the first study in Rwanda and one of very few studies in the broader East African region analyzing the microbial assemblages present in communities of field-collected mosquitoes (Osei-Poku et al., 2012; Y. Wang et al., 2011b). The two sites studied here provide a broader idea of mosquito and microbial community assemblages in both disturbed, human-occupied habitat (Nyabugogo), and preserved, natural second-growth forest habitat (Huye Campus). To best understand how to control arboviral and mosquito-borne parasite transmission using microbial control methods, it is essential to understand both domestic and sylvatic mosquito systems (Table 2).

**Table 2.**
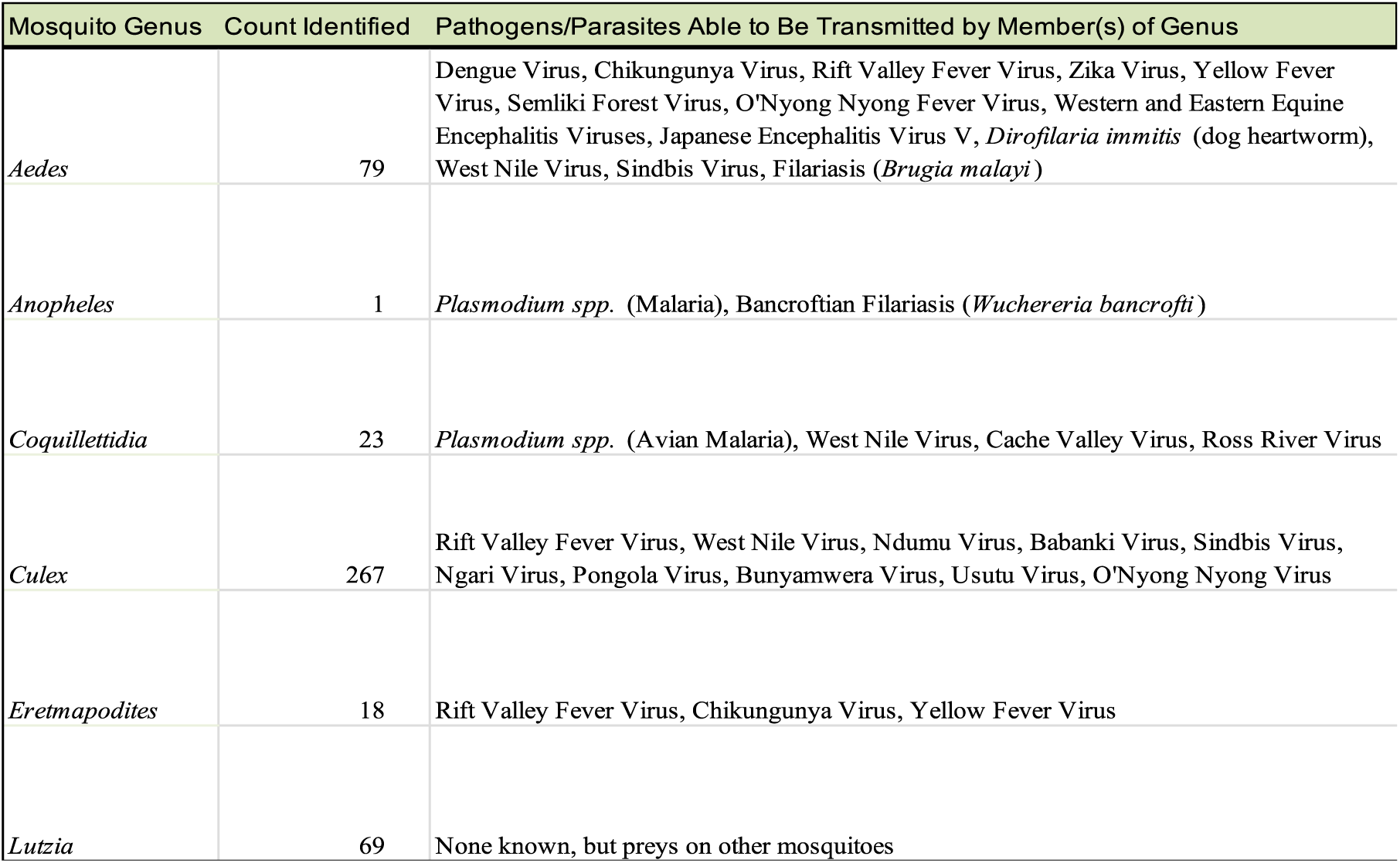
Count of mosquito genera positively identified in samples collected in this study in Rwanda and classified using COI sequencing. Pathogens and parasites that each genus is capable of transmitting are listed. No viruses were directly characterized as being present in any of the collected samples and this list is not intended to be exhaustive. (Ajamma et al., 2018; Appawu et al., 2000; Arum et al., 2015; Becker et al., 2010; Benelli et al., 2015; Biteye et al., 2019; Cansado-Utrilla et al., 2019; CG et al., 1998; Corbet et al., 1961; Culicidae Classification | Mosquito Taxonomic Inventory, n.d.; Devi et al., 2018; Diallo et al., 2011; Dohm et al., 1995; Faran et al., 1987; Fontenille et al., 1998; Fontenille & Jupp, 1989; A. D. Haddow et al., 2016; A. J. Haddow, 1946; Hamer et al., 2008; Hylton, 1969; Jeffery et al., 2002; Jupp & McIntosh, 1970a, 1970b; Karungu et al., 2019; Kilpatrick et al., 2005; Kovendan et al., 2012; LaBeaud et al., 2011; Lutomiah, Omondi, et al., 2014; Lutomiah, Ongus, et al., 2014; Mangiafico, 1971; McGreevy et al., 1974; Moncayo et al., 2000; Musa et al., 2020; Mutebi et al., 2013b; Ndiaye et al., 2016b, 2016c; Nikolay et al., 2012; Njabo et al., 2009a, 2009b; Pullan et al., 2010; Sardelis et al., 2001; Sharma et al., 2005; SMITHBURN et al., 1949; Snow & Boreham, 1978; Traore-Lamizana et al., 1994; Valkiunas et al., 2008; Waddell et al., 2019)

In terms of alpha diversity metrics (Observed sOTU Richness and Shannon Diversity Index), we did not see any strong trends or differences between groups, including genera, species, and sampling location. We hypothesize that this lack of differences between genera, species, and locations may be due to similar colonizable area and selection pressures in the internal environment of the mosquitoes, though further work must be undertaken to address this. Differences in alpha diversity between visibly bloodfed and non-bloodfed mosquitoes were anticipated, based on prior studies regarding the reducing environment in the mosquito gut following bloodfeeding by females. However, our data showed a weak difference between these groups (Y. Wang et al., 2011a). This is likely due to the visual nature of confirming bloodfed status of females in this study. Only female mosquitoes that were visibly engorged upon capture were considered bloodfed, which does not include females that had previously fed but digested the blood and were no longer engorged.

In terms of beta diversity metrics (Jaccard and Weighted Unifrac distances), several interesting trends emerged in the data. Genera, species, and location were all highly significantly different for both Jaccard and Weighted Unifrac comparisons, demonstrating that taxa and broader habitat play a key role in the community assemblage of the mosquito microbiome under field conditions. Bloodfed status, catch method, and mosquito sex were all non-significantly different. The lack of difference between catch methods indicated that there was not a sampling bias based on the method of capture utilized for mosquitoes. The absence of difference in beta diversity for bloodfed status indicates that, similarly to our alpha diversity measures, recently fed females that had digested blood were probable in the non-bloodfed group. Mosquito sexes were not different in terms of microbial community assemblage, though they were anticipated to be different based on differences in feeding between sexes. Here, we hypothesize that environmental drivers and mosquito microhabitat use are more important than sex in terms of the formation of internal microbial communities, though this will also require additional investigation to better understand these dynamics.

The core microbiome, or the 20 most abundant sOTUs appearing in nearly all samples in high relative abundance, included numerous members well documented as being abundant in mosquito microbiomes in other studies around the world (Berhanu et al., 2019; Buck et al., 2016; Dennison et al., 2014; Dickson et al., 2017; Novakova et al., 2017; Osei-Poku et al., 2012; Y. Wang et al., 2011a). *Asaia, Serratia*, and *Wolbachia* are genera of bacteria of particular interest for controlling arboviral transmission, and all three were well represented in the core microbiomes of the mosquitoes collected in this study. The average number of sequences (from data rarefied at 2000 sequences/sample) of these genera was much higher at the Nyabugogo site than the Huye Campus site. The distribution of mosquitoes carrying these microbes was not continuous across any given genus, but rather varied greatly by species. Several mosquitoes that appeared in abundance at both sites had much higher average sequences of these bacteria at Nyabugogo than at the Huye Campus. While these interactions are inherently complex, we suggest that the likelihood of colonization by these microbes of interest may be an interaction of predisposition for colonization by mosquito taxa and microhabitat use.

In order to more fully understand mosquito microbial interactions *in situ*, studies further examining the level of influence that specific environmental factors, such as temperature and rainfall, have over the colonization of mosquitoes. These studies should be undertaken both in a laboratory and field setting, as controlling variables in the field can present challenges, whereas work in the lab does not fully realize all potential variables influencing colonization of the mosquito. We would also recommend pursuing additional studies on the eukaryotic, fungal, and viral members of the mosquito microbiome for a more complete picture of these microbial communities in a field setting.

Our findings help to expand on our understanding of wild mosquito microbiomes in an area of the world that is particularly at risk for high arboviral transmission. In order to safely utilize microbial tools for mosquito control, such as *Wolbachia*-infected mosquito releases, we must first understand the natural communities of both mosquito hosts and microbial inhabitants and the factors that influence these interactions.

## Author Contributions

ATP and DCW conceived, planned, and acquired funding for this study. ATP, JDN, MK, SM, IWT, SGL, and JDJ carried out the experiments. ATP analyzed the data. ATP wrote the manuscript with input from the other authors.

## Acknowledgements

We would like to thank our collaborators at the University of Rwanda and the Center of Excellence in Biodiversity. Content of this manuscript was previously included in Amanda Tokash-Peters’ dissertation.

## Data Availability

The datasets generated and analyzed during the current study are publicly available. This data can be found here: http://www.ncbi.nlm.nih.gov/bioproject/849234.

## Supplementary Figures

**S1.**
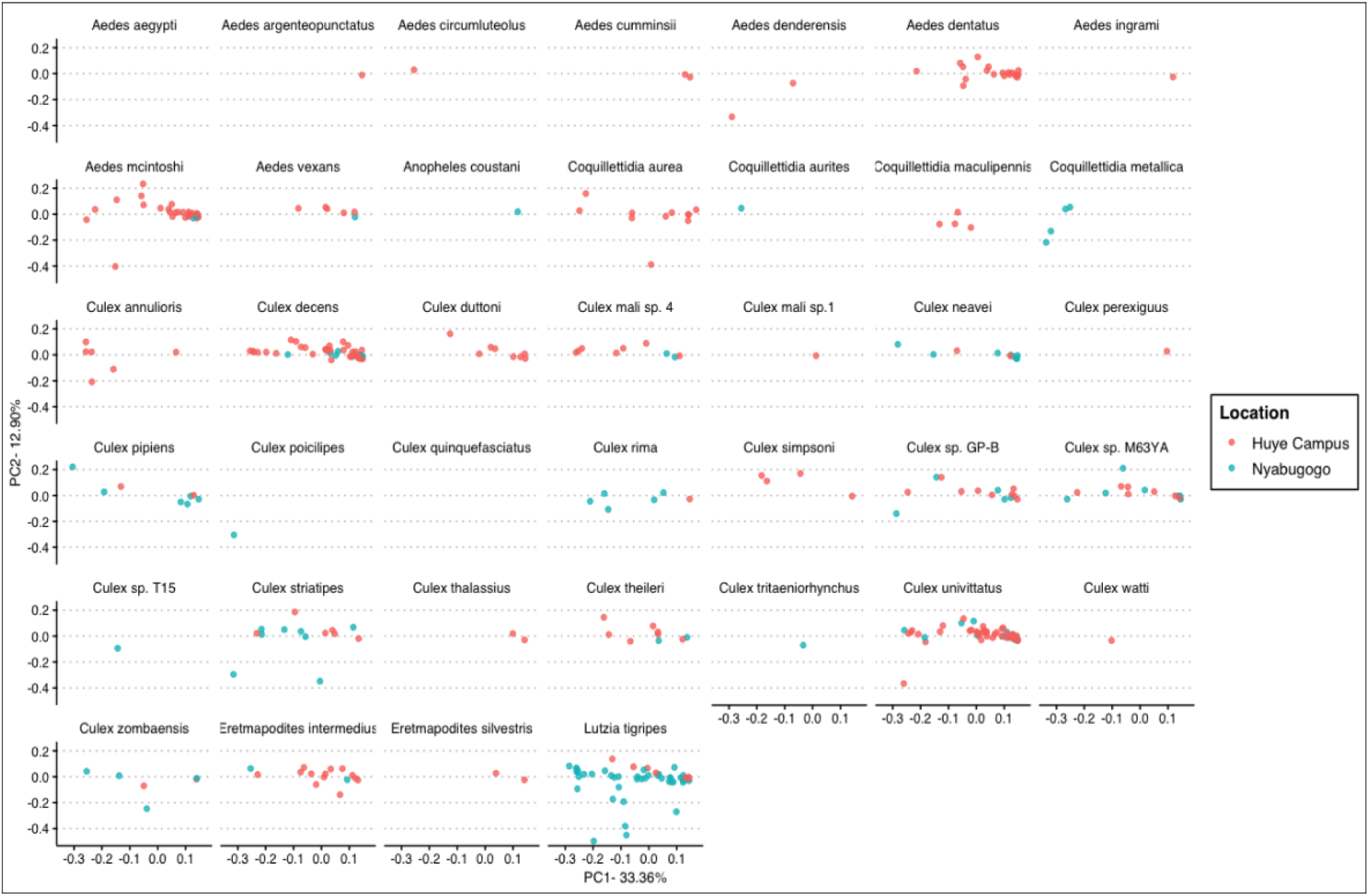
Principle coordinate analysis made with weighted Unifrac distances to demonstrate the relative community composition of the microbiome for each species by sampling location. Some species were only found at one location and therefore do not show both sites in the figure.

